# BiGen: Integrative Clinical and Brain-Imaging Genetics Analysis Using Structural Equation Model

**DOI:** 10.1101/2020.02.04.934596

**Authors:** Samar S. M. Elsheikh, Emile R. Chimusa, Alessandro Crimi, Nicola J. Mulder

## Abstract

The identification of genetic variants associated with complex brain diseases has evolved in the past decades. Studies in the field have taken different approaches and study designs including genome-wide association studies. Neuroimaging and connectomics have also improved our understanding of the structural connectivity of the human brain and produced reliable measurements. Combining both neuroimaging and genetic characteristics significantly contributes to understand their complex relationship in affecting behaviour and cognition. Throughout this thesis we proposed analysis pipeline to study the association between imaging and genetics of two different types of brain disease, which is, Alzheimer’s disease and glioblastoma. We observe the need for a unified model to study the complex interplay between genetic, environmental and clinical, neuroimaging and phenotype features. In this chapter, we developed BiGen, a mathematical model to measure the inter-correlation structure through the integration of genetic, environmental, neuroimaging and disease measurements. We utilised the structural equation model and used a path construct of latent variables to study the hidden association between genes and brain-related diseases, mediated by connectivity characteristics. We applied BiGen to simulated data and to a dataset from the Alzheimer’s Disease Neuroimaging Initiative.

## 1. Introduction

Imaging genetics is a rapidly growing field that focuses on the identification of genetic variants associated with complex brain diseases. In doing so, imaging genetics combines brain imaging technologies outputs with genetic data. Depending on the hypothesis of interest and data availability, imaging genetics studies can focus on 1) the association between each location in the brain, namely voxel, with each single nucleotide polymorphism (SNP), e.g. Voxelwise Genome-Wide Association Study (vGWAS) (Shen et al. 2010; Stein et al. 2010), 2) the relationship between a single phenotype, and, such as the hippocampus volume, such studies are called candidate phenotype studies (Stein et al. 2012), or 3) Other studies consider multiple imaging endophenotype associations with candidate genotypes.

Recently, brain connectivity and other magnetic resonance imaging (MRI) features have been used as phenotypes in GWAS and other omic studies. Brain connectivity metrics are derived from the connectome - a representation of the brain as a single network extracted from the diffusion tensor imaging (DTI) (Sporns et al. 2005). The nodes of a connectome represent distinct regions in the brain and the links are the water tracts connecting each pair of regions. Some studies used GWAS with connectivity metrics to study the healthy brain (JahanshAD et al. 2013), Alzheimer’s Disease (AD) (Elsheikh et al. 2020), and other brain diseases (Alloza et al. 2018; Thompson et al. 2010). Imaging genetics uses a variety of study designs that integrate specific genetic information (Thompson et al. 2010), Elsheikh et al. (2019), for example, studied the effect of targeted gene expression on local and global connectivity features in AD.

The recent imaging genetics literature has proposed multivariate methods to understand the influence of multidimensional genotypes on multi-phenotype imaging characteristics. As summarised by Liu and Calhoun (2014); the multivariate methods in the field of imaging genetics need to consider three factors; 1) the dimensionality of genotypes and intermediate neuroimaging phenotypes, 2) the importance of managing the confounding factors in order to reveal the imaging phenotypic effects, 3) the population structure of the sample under study. Considering these criteria, previous studies proposed multivariate analysis methods utilising sparse reduced-rank regression (Vounou et al. 2010), sparse partial least square (Le Floch et al. 2012), parallel independent and principal component analysis (Liu et al. 2009) and sparse canonical correlation analysis (Chi et al. 2013). Other methods considered longitudinal multivariate imaging genetics pipelines to study the relationship of gene expression and genome-wide variants with brain connectivity metrics (Elsheikh et al. 2018, 2020, 2019; Lu et al. 2017).

The structural equation model (SEM) (Bollen and Long 1993) is a multivariate technique that studies the complex causal relationship between a set of endogenous and exogenous latent constructs through measured or observed variables. SEM estimates the structural relationship between the latent variables using a set of multiple regression models and factor analysis. Recently, SEM has been applied in the context of mapping genotype and phenotype in complex diseases and imaging genetics. Specifically, SEM is a promising tool in merging gene regularity networks with post-GWAS approaches to improve the understanding of cell signaling, and metabolic pathway translation into phenotypes (Nuzhdin et al. 2012). Grotzinger et al. (2018) proposed Genomic SEM, a tool that identifies between-diseases similarity and genetic architecture, with an application to psychiatric disorders. Huisman et al. (2018) applied the SEM to understand the spatial change within brain regions affected by gene expression using a dataset from Alzheimer’s Disease Neuroimaging Initiative (ADNI). They used healthy brain information from spatial transcriptome datasets to identify the model construct.

Here, we propose a method to identify the causal effect of genetics on multiple phenotypes, including neuroimaging and disease measures. We utilised the structural equation model to estimate the association between various latent constructs, including disease, endophenotypes, covariants and genetics. In our model, the latent variables were estimated using observed measurements and the relationship between disease and genetics was estimated accounting for the intermediate effect of neuroimaging endophenotype. The latent genetic variable was measured through observed gene expression, considering the interaction of proteins, while the disease and endophenotype latent variables were measured through the Clinical Dementia Ratings (CDR) and global connectivity metrics, respectively. Moreover, we controlled for confounding factors through a separate latent construct, inferred through some environmental factors. We applied the new tool to simulated data and a dataset from ADNI.

## 2. Materials and Methods

In this section we describe our proposed SEM model, BiGen. The model has two parts, the structural model and the measurement model which are described in the following sub-sections. Thereafter we discuss the estimation of these models.

### Structural Model

The structural model in a SEM relates the exogenous and endogenous latent variables (construct) and studies their associations. The advantage of using latent variables is to control for the measurement errors on the overall SEM. In BiGen, the structural model consists of four latent variables, these are genetics, confounders, endophenotypes and disease. These four latent constructs are measured through some genetic measurements, environmental and other risk factors, neuroimaging characteristics and disease measures of interest. See Figure 1 where latent variables are shown as circles.

**Figure 1.**
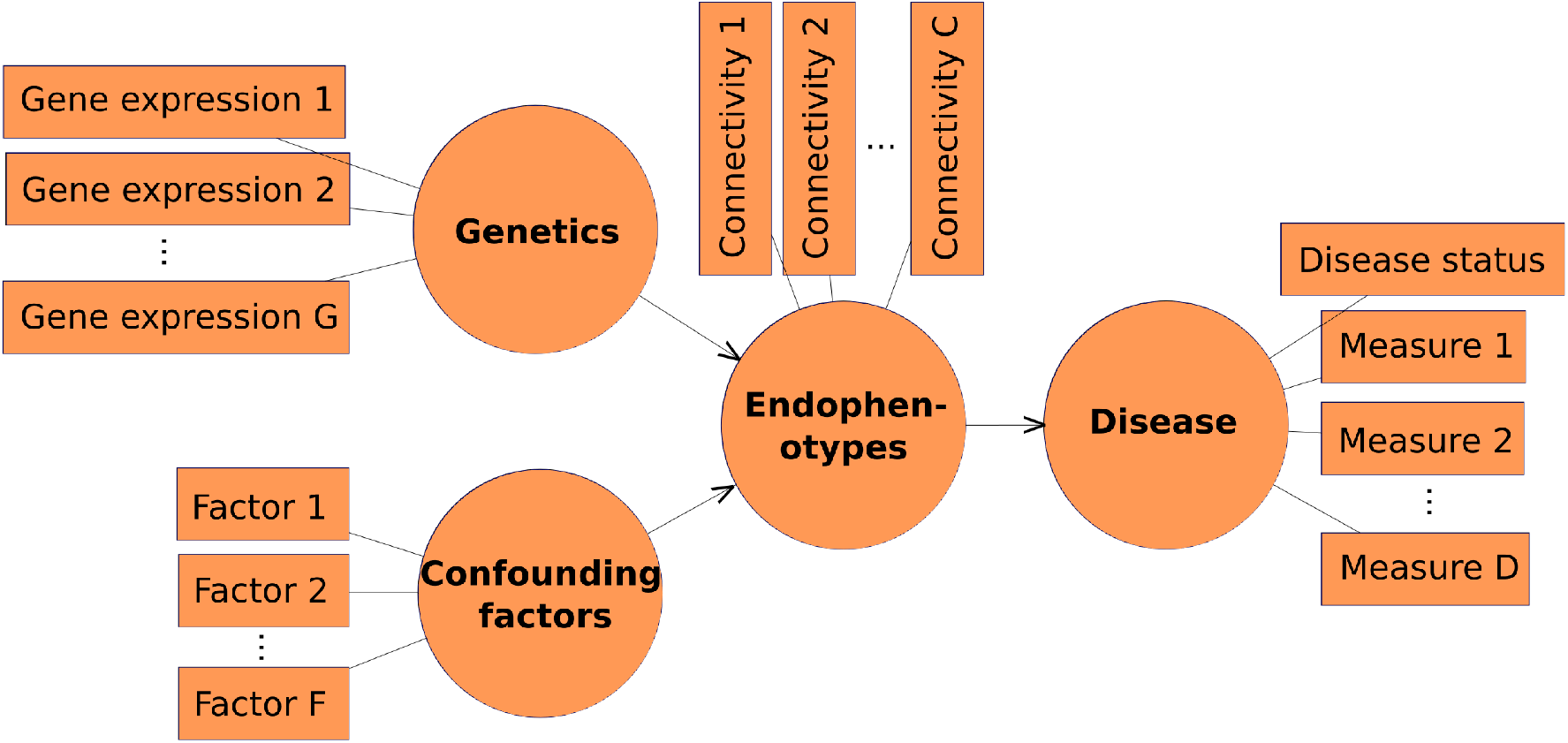
The BiGen model showing the latent variables (in circles) and the observed measurements (rectangles). The structural model shows latent variables and the relationship between them, while the measurement models connects the measurement variables with latent constructs.

### Measurement Model

The measurement model focuses on the relationship between latent variables and their observed measurements (indicators). The measurements are shown as rectangles in Figure 1. Specifically, in BiGen, the genetics and confounding factors constructs relate to the endophenotypes. The endophenotypes have one direct relationship with the disease construct. Moreover, the endophenotypes mediate between genetic (and confounding factors) and the disease construct. This mediation is considered to account for the indirect effects of genetic factors on the disease through the endophenotypes. BiGen analyses the effect of genetic-endophenotype interaction on the disease.

### Model Estimation

We applied the BiGen model to an AD dataset from ADNI. In this case, the constructs were measured and inferred through a set of observed measurements. Specifically, the genetics construct was measured through gene expression values that interact with one another. These gene expression values correlate the measurements of other latent variables with the genetics construct. The confounding factors were basically the random effects or other risk factors that are not genetics. Here, other risk factors, if available, could be used as measurements for the latter construct, such as blood pressure, cholesterol level and heart diseases. A set of global connectivity metrics were used to measure the endophenotypes construct. In this work we used the global connectivity metrics, namely, global efficiency, Louvain modularity, transitivity and characteristics path length. Finally, the disease construct was measured through a set of clinical dementia ratings. Section 2. and Section 2. explain the simulated and AD datasets in more detail, respectively.

The structural model consists of two exogenous constructs (Figure 1), these are genetics and confounding factors, and two endogenous constructs, endophenotypes and disease. To estimate the SEM we followed two estimation steps. In the first step, we estimated the latent variable scores through an iterative algorithm that does not assume any distribution of the measurements or constructs. In doing so, we adjusted the partial least squares (PLS) SEM iterative algorithm proposed by Lohmöller (1989). The approach estimates the latent scores in four iterative steps until a stop criteria is met. The second step estimates the path coefficients and outer weights through ordinary least square models. This basic PLS algorithm has been applied successfully to many fields, including advertising research (Henseler et al. 2012). In our model, we used non-parametric methods and a summary of the steps we adopted is provided below.

In our first step, we followed a set of sub-steps that aim to estimate the latent scores iteratively until convergence. More specifically, considering the notation shown in Figure 2, the first sub-step is to estimate *w_i_j* for all the measurement models connecting *x_i_j* with *y_j_*. Initially, we initialised these weights to 1, and updated them iteratively. The second sub-step is to estimate the inner weights (*b_ik_*; see Figure 2). This is followed by an approximation of the latent variables 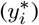, and in the last sub-step we estimate the outer weights. These four sub-steps are summarised below. The tolerance is calculated at the end of each iteration, and this is simply the difference between the current outer weight and the previous one. The threshold here was set to 1*e* − 6.

Step 1.1 External approximation of latent variables, 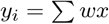
Step 1.2 Estimation of inner weights, *b_ik_* = *ρ_ik_*, then normalize.
Step 1.3 Internal approximation of latent variables, 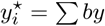
Step 1.4 Estimation of outer weights, *w* = *ρ_xy^⋆^_* (where *ρ* is Spearman correlation coefficient), then normalize.

**Figure 2.**
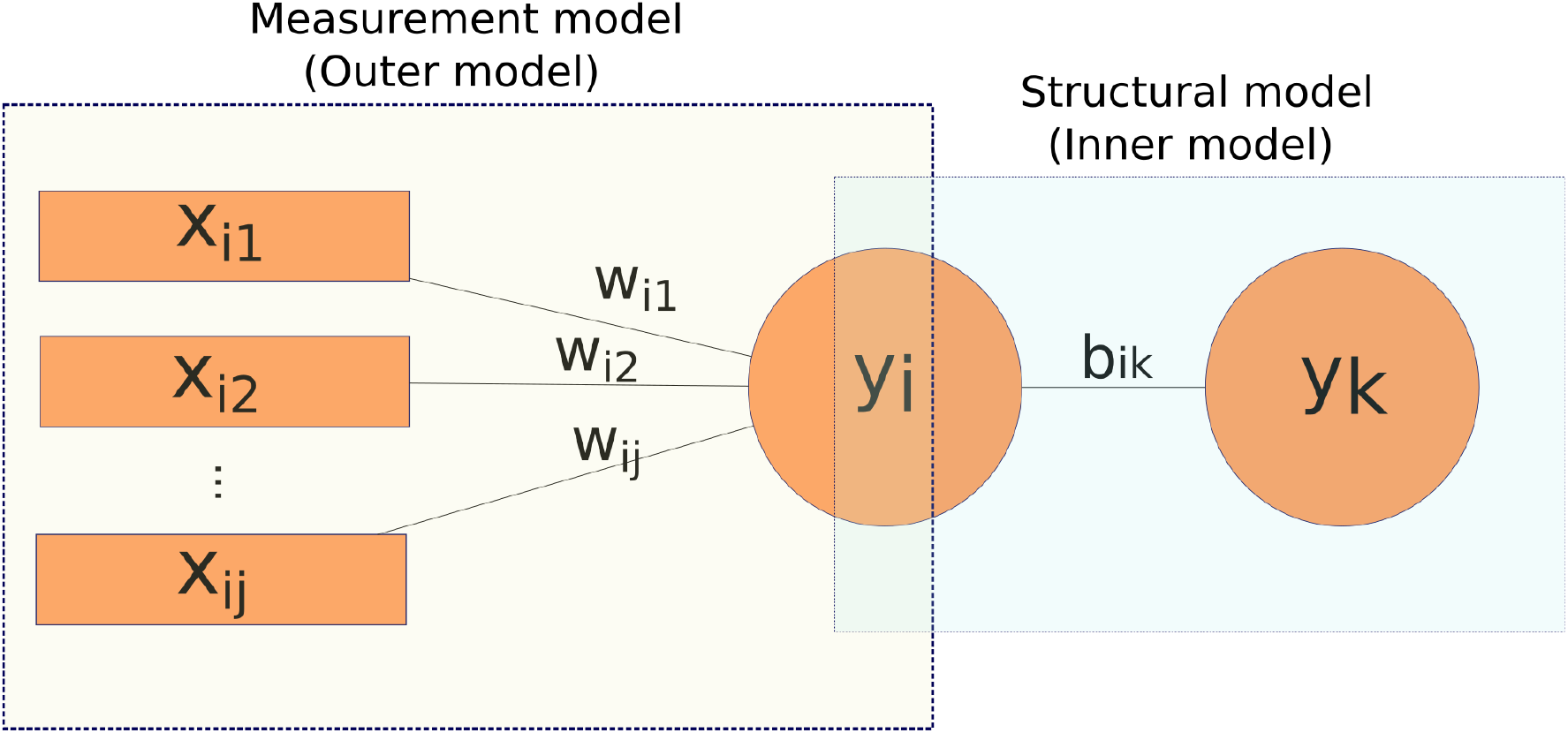
Notation used in model estimation. The left box represents the measurement model, and the right box represents the structural model. The outer weights (*w*_*i*1_, *w*_*i*2_ … *w_ij_*) connect the indicators (*x*_*i*1_, *x*_*i*2_ … *x_ij_*) to the latent variable *y_i_*. The inner weight (*b_ik_*) associates the latent variables in the structural model (*y_i_* and *y_k_*).

As a second step, BiGen estimates the final path coefficients and outer weights through a (quantile) regression model. Besides the inner model shown in Figure 1, BiGen calculates the inner of the interaction terms: genetics×endophenotype and confounding factors×endophenotype and estimates their effect on the disease.

### Simulated Data

We simulated a dataset to test BiGen and compare it to the PLS-SEM. Firstly, we randomly generated a single continuous measurement variable of the disease construct. We then created another two measurement variables with strong correlation with the first measure, but with some randomness. We simulated three connectivity variables that are associated to one another, and to the three measures of the disease measurements, with some randomness. Similarly, we simulated the genetic and confounding factor measures (three for each).

To simulate the variables, we used a random range of parameters. Specifically, we simulated 12 variables as described above, each follow the random distribution with different mean (ranges between 100 and 300) and standard deviation (ranges between 2 and 18). Figure 3 shows the distribution of the simulated data, while the pattern of association between the variables is shown in Figure 4. The sample size ofthe simulated data is 5000.

**Figure 3.**
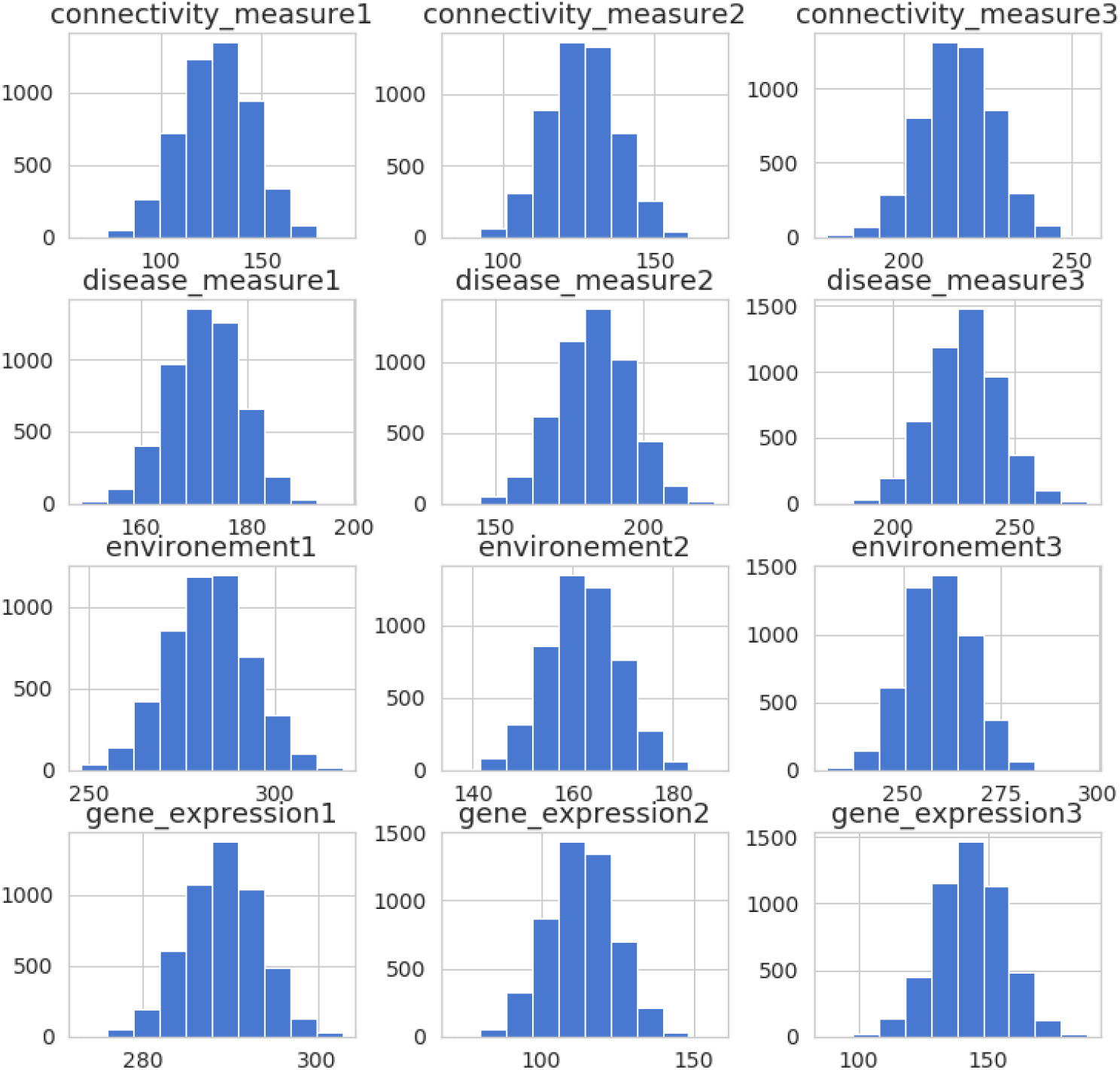
The distribution of all the simulated measurement variables used to fit the proposed BiGen model.

**Figure 4.**
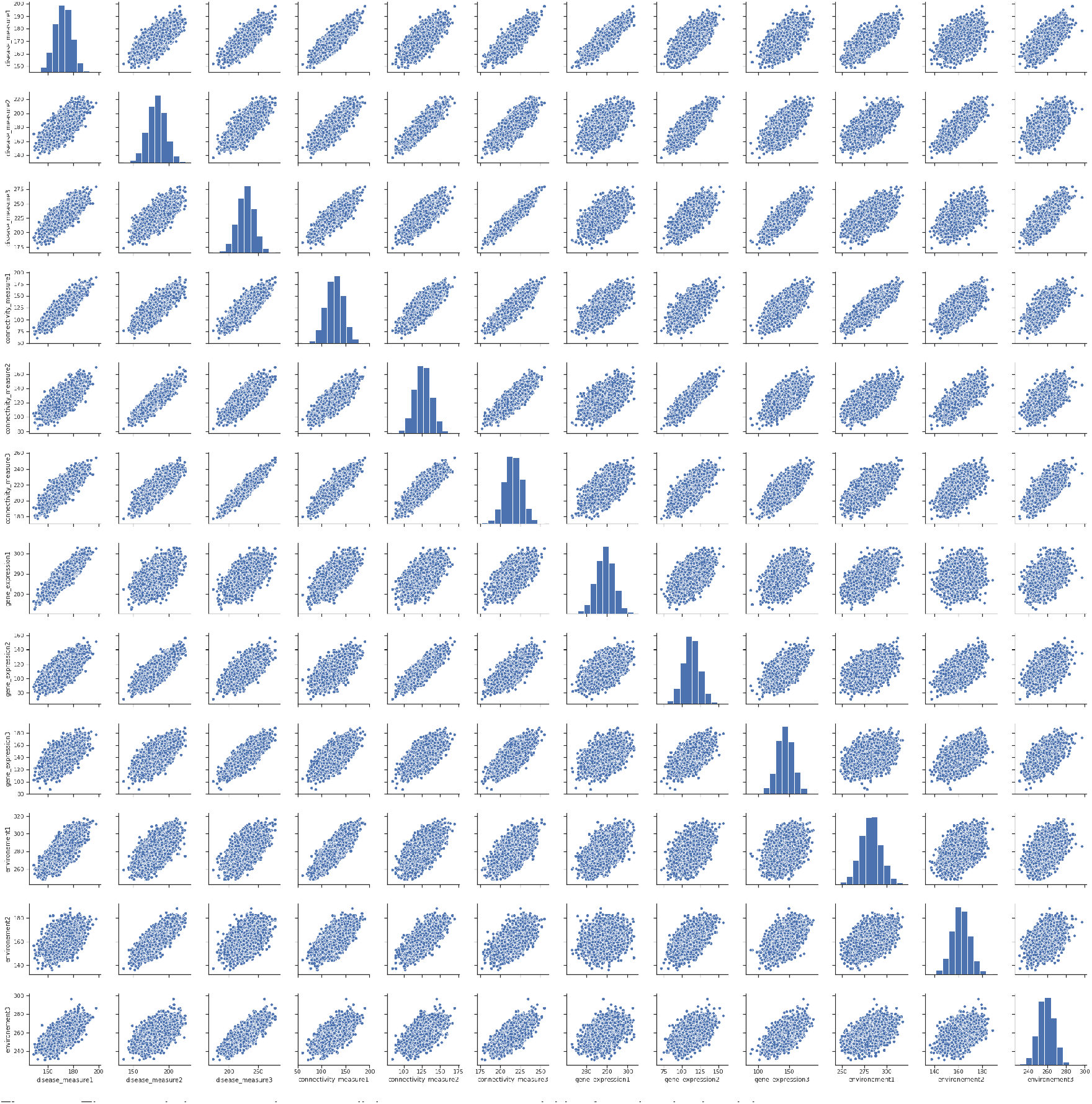
The association pattern between all the measurement variables from the simulated data.

### Alzheimer’s Disease Dataset

We also applied our model to an AD dataset from ADNI (available at adni.loni.usc.edu). Specifically, we used the information of ADNI participants as measurement variables to fit our proposed model (as shown in Figure 1). For the 1) disease, 2) endophenotypes, 3) confounding factors and 4) genetics we used the following measurements; 1) three CDR measures, namely, memory, judgment and problem solving, and home and hobbies scores, 2) the global connectivity metrics, namely, Louvain modularity, transitivity, global efficiency and characteristics path length (all explained in Section ??), 3) the education level and gender of participants, and 4) the expression of a set of genes that interact with one another, namely *APP, SORL1, ADAM10, ApoE* and *PSEN1*. We obtained the expression values following the same steps explained in ??. The protein-protein interaction data were derived from STRING database (v11.0), accessed at https://string-db.org (Szklarczyk et al. 2014, 2018). We used the absolute difference between the baseline and follow-up visits for the disease and endophenotypes measurements. Figure 5 shows the distribution of all pre-described measurements, while Figure 6 shows their association patterns. In total, we managed to obtain 46 samples.

**Figure 5.**
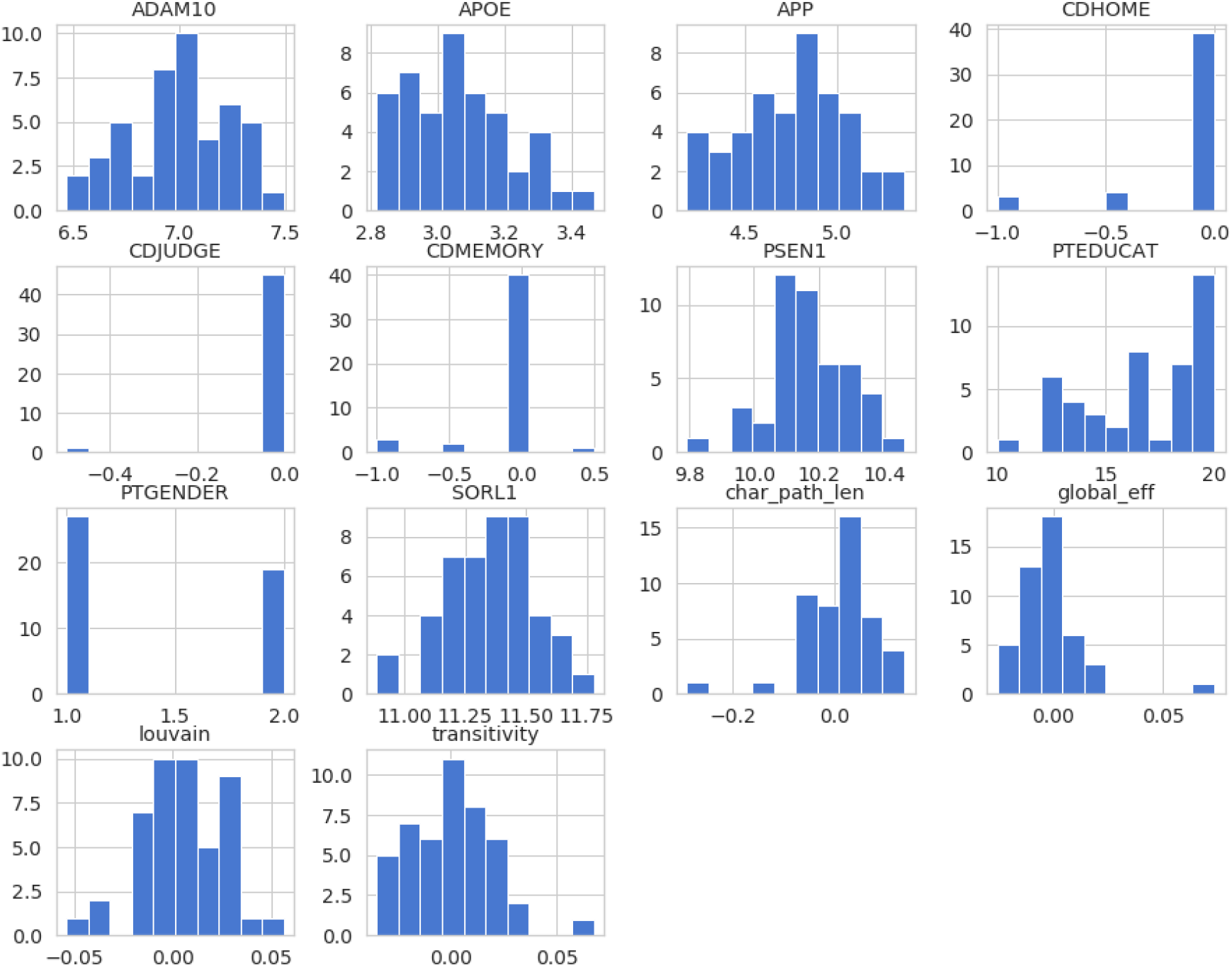
The distribution of all the measurement variables obtained from ADNI dataset and used to fit the proposed BiGen model.

**Figure 6.**
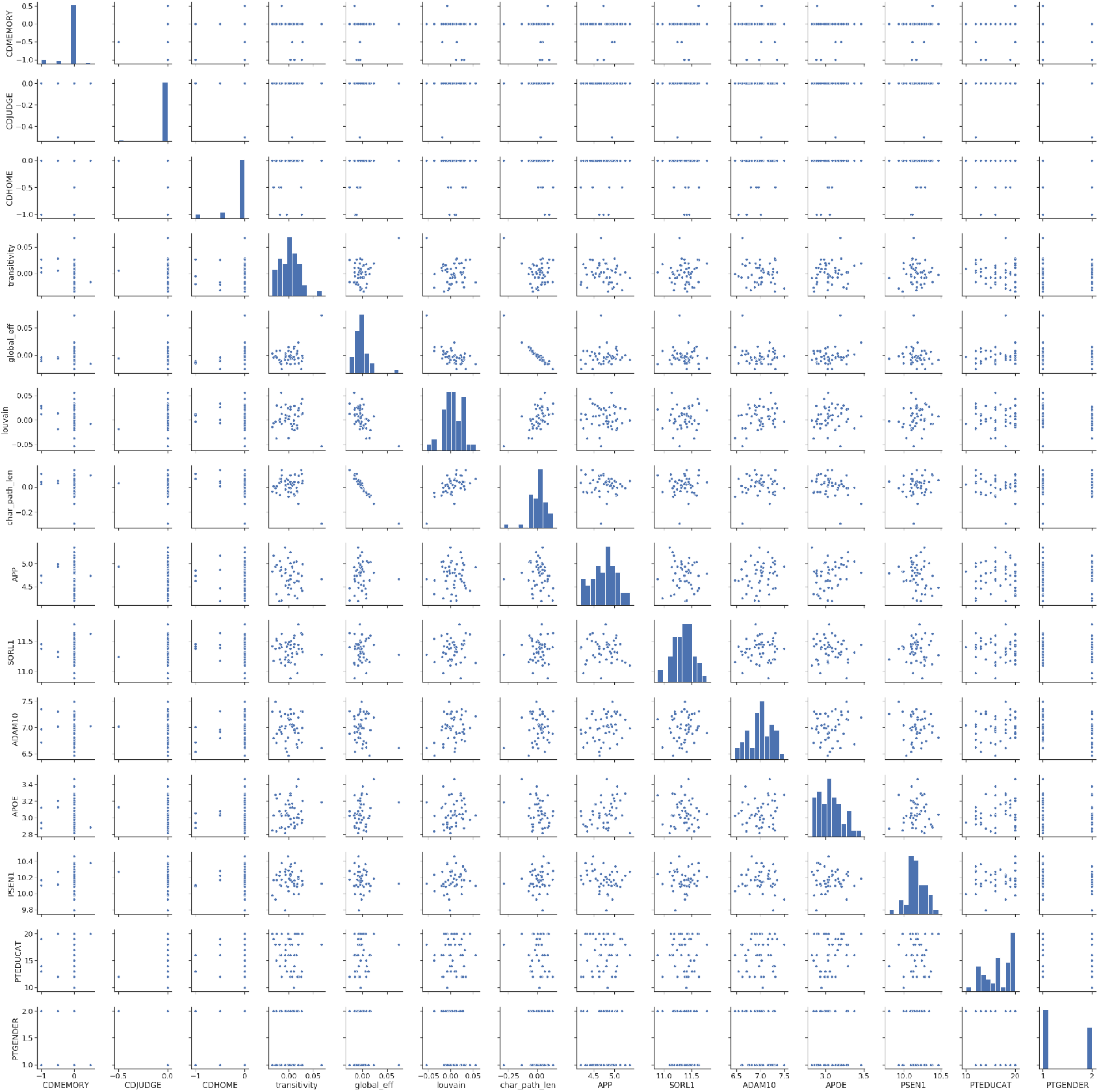
The association patterns between all the measurement variables obtained from ADNI dataset.

#### Software

To conduct the analysis described here, we used Python 3.7.1 and made our scripts available under the MIT License and accessible at: https://github.com/elssam/BiGen.

## 3. Results

### Simulation

#### PLS-SEM

Using the simulated measurement variables described in Section 2. we fitted, firstly, the PLS-SEM, the path coefficient and inner weights are both shown in matrices 1. While the interaction terms (described in Section 2.) inner weights are shown in matrix 2. We then fitted BiGen (as described in 2.) to the same data and obtained the path and inner weights that are shown in matrices 3, similarly, we obtained the inner weights estimates for the interactions, this is shown in matrix 4.

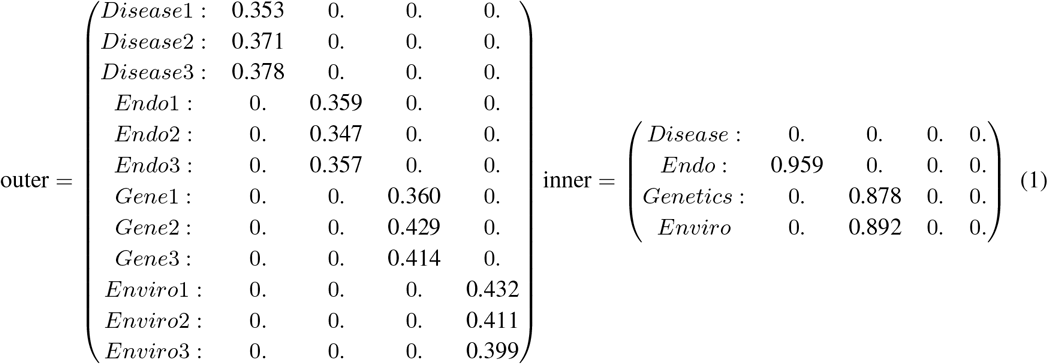

#### PLS-SEM wtih Interaction

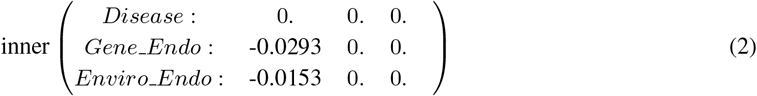

#### BiGen

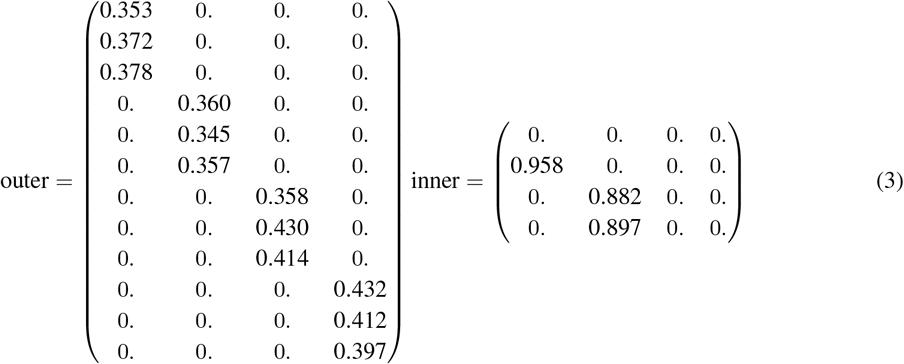

#### BiGen wtih Interaction

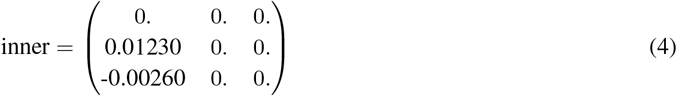

### Application to ADNI

Then, we applied both BiGen and SEM-PLS to the AD measurement variables discussed in Section 2.. Similarly, we considered the interaction model in both applications. The path coefficient and inner weights obtained form applying the PLS-SEM are both shown in matrices 5, while the interaction inner weights are shown in matrix 6. We then fitted BiGen to the AD dataset and the results of the path and inner weights are shown in matrices 7. We finally computed the inner weights estimates for the interactions (see matrix 8).

Generally, we observe that applying BiGen and PLS-SEM to the simulated data gave the same results (compare: matrices 1 vs matrices 3, and matrix 2 vs matrix 4), however, in the AD application the two models gave slightly, though not significantly different results (compare: matrices 5 vs matrices 7, and matrix 6 vs matrix 8). We observe that BiGen produced different effects of the inner path (inner matrix 7) compared to PLS-SEM (inner matrix 5) when applied to ADNI dataset. The overall effect of the genetic and confounding factor latent constructs on the endophenotype are 0.113 and 0.216 in BiGen, while they are −0.337 and −0.150 in PLS-SEM. Though non of these were statistically significant, we observe that the contribution of genetic factors on the endophenotype are lower in BiGen, the positive sign indicates the direction of the relationship. This might indicate that PLS-SEM overestimates the inner path effect of genetic construct on the endophenotype and underestimates the confounding factor effect on the endophenotype.

#### PLS-SEM

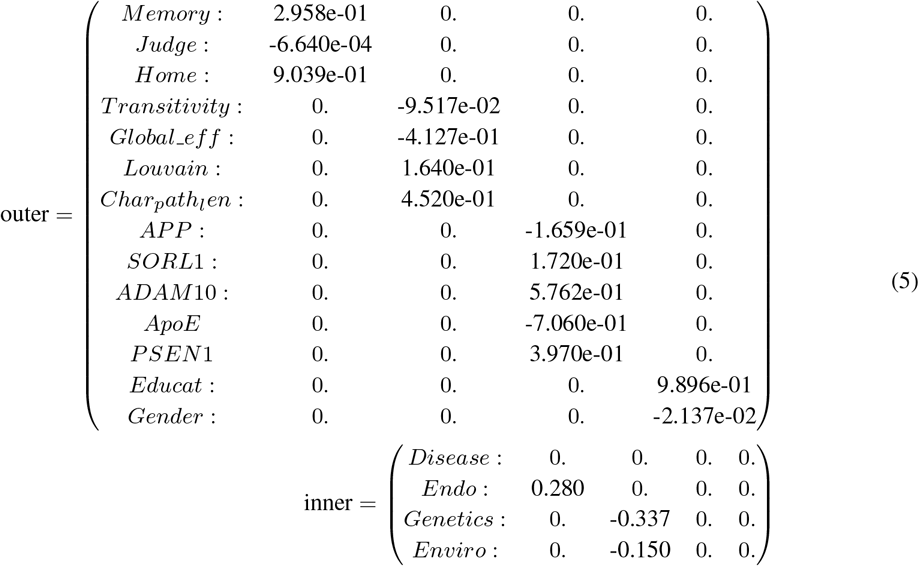

#### PLS-SEM wtih Interaction

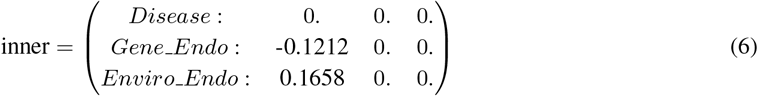

#### BiGen

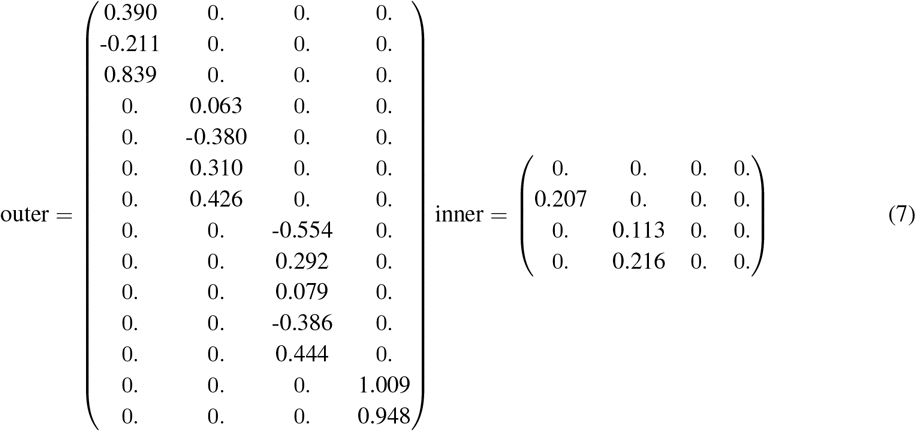

Similarly, the interaction terms were smaller when we fit BiGen to the ADNI dataset compared to PLS-SEM (compare the BiGen inner interaction terms in matrix 8 vs SEM-PLS inner interaction terms in matrix 6). Our overall explanation to these results, including the simulation results, is that the simulated data are made to be symmetric and normally distributed, therefore, both the PLS-SEM and BiGen models offer a similar performance, while in the ADNI application, the pattern of distribution is not symmetric and the data types are different, which is always the case in real life applications. However, we believe that more samples are needed to better understand the performance of BiGen in the ADNI dataset.

Table 1 shows a brief comparison of the PLS-SEM and BiGen results in the application to simulation and ADNI datasets. We note that both BiGen and PLS-SEM were fast (see the Time column), even though the ADNI application needed more iterations in both models. To get the interaction terms, the same calculations in the original model are performed, except that we need to estimate a different path coefficients in the second step (see Section 2.).

**Table 1.** SEM, BiGen with and without interaction results for simulated and AD dataset.

| Application | Simulation |  |  | AD |  |  |
| --- | --- | --- | --- | --- | --- | --- |
|  | Iterations | Tolerance | Time* | Iterations | Tolerance | Time |
| (1)2-7 |  |  |  |  |  |  |
| SEM | 2 | $7.918e-08$ | 202 ms<sup>+</sup> | 23 | $6.702e-07$ | 241 ms |
| SEM (interaction) | 2 | $7.918e-08$ | 189 ms | 23 | $6.702e-07$ | 184 ms |
| BiGen | 2 | $1.367e-07$ | 502 ms | 26 | 0.000 | 1.32 s<sup>†</sup> |
| BiGen (Interaction) | 2 | $1.367e-07$ | 521 s | 26 | 0.000 | 1.26 s |

\* Running time
<sup>+</sup> Millisecond
<sup>†</sup> Second

#### BiGen with Interaction

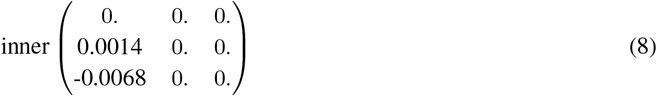

The main difference between the above results and what was done in Chapter ?? and Chapter ?? is that here we did not explicitly test the pairwise associations of the observed measurement variables (e.g, global connectivity vs gene expression), but rather, we tested their combined association through the latent constructs (e.g. the effect of genetic on the endophenotype), this was finally estimated by the inner weights, shown in the matrices above. For example, Table ?? shows that global connectivity metrics associated with the expression of *APP, ADAM10, ApoE* and *PSEN1* with correlation coefficients −0.2602 (characteristic path length), −0.236 (Louvain modularity), −0.25 (characteristic path length) and −0.299 (transitivity). On the other hand, BiGen can not measure the association in the same way, but rather, 1) we observe the contribution of the previous gene expression on the genetic construct, these are −0.554, 0.079, −0.386, 0.444, respectively (see outer matrix 7), 2) BiGen also allows measuring the effect of the genetic construct on the endophenotype, and that was 0.113 (see inner matrix 7).

In Chapter ??, we tried to test different hypotheses individually, we tested the effect of gene expression on the connectivity metric using both quantile regression and Spearman association coefficient, we then examined the effect of brain connectivity on the CDR measurements, after which we tested their combined effect on the CDR measurements using ridge regression. BiGen allows us to test all these hypotheses (including the interaction terms between them) simultaneously in only two steps and with less computational complexity. Additionally, in Chapter ?? we conducted a GWAS on the ADNI dataset and integrated the summary statistics obtained from GWAS at a gene-wide level, we found that some genes were significantly associated with global connectivity metric (e.g *ANTXR2, IGF1* and *OR5L1*). However, to study the effect of these genes using BiGen, we would consider the protein-protein interactions, and incorporate the other factors shown in Figure 1.

## 4. Conclusion

The SEM is a commonly used multivariate technique in many scientific fields, it studies the structural relationships between variables that are hard to measure directly. Analysing imaging genetics data needs not only a multivariate technique, but also one that considers the complexity, heterogeneity and multicollinearity of both the imaging and genetics parts. Here we propose BiGen, a Python tool that analyses the structural relationship between four constructs simultaneously, these are, genetics and disease, mediated by the neuroimaging endophenotypes and considering some environmental or confounding factors. All the four latent constructs are measured through observed indicators and the model is specifically made for brain-related disease, with an application to Alzheimer’s disease through ADNI data. BiGen adjusts the PLS-SEM by using non-parametric regression and association tests in estimating the path model and latent variable scores, respectively. This causal predictive model is flexible, simple and computationally inexpensive. Additionally, BiGen has a satisfactory performance in small samples and assumes no distribution or hypothesis of the data.

Our model can easily be extended to include more indicators and constructs. For example, one can add more imaging phenotypes, or rather, incorporate the local connectivity metrics to the endophenotype latent construct. Moreover, the model is flexible to more complexity in the path model, this is especially useful when studying the direct relationship between genetics and disease phenotypes, or the gene-environment interaction effects on the disease or endphenotypes. Over all, we propose that BiGen is suitable for unvealing the complex interplay between neuroimaging and genetics in the context of brain-related diseases.

